# Release of insulin granules by simultaneous, high-speed correlative SICM-FCM

**DOI:** 10.1101/2020.09.17.302372

**Authors:** Joanna Bednarska, Pavel Novak, Yuri Korchev, Patrik Rorsman, Andrei I. Tarasov, Andrew Shevchuk

**Author notes:** Correspondence to: AS and AIT.

## Abstract

Exocytosis of peptides and steroids stored in a dense core vesicular (DCV) form is the final step of every secretory pathway, indispensable for the function of nervous, endocrine and immune systems. The lack of live imaging techniques capable of direct, label-free visualisation of DCV release makes many aspects of the exocytotic process inaccessible to investigation. We describe the application of correlative scanning ion conductance and fluorescence confocal microscopy (SICM-FCM) to study the exocytosis of individual granules of insulin from the top, non-adherent, surface of pancreatic β-cells. Using SICM-FCM, we were first to directly follow the topographical changes associated with physiologically-induced release of insulin DCVs. This allowed us to report the kinetics of the full fusion of the insulin vesicle as well as the subsequent solubilisation of the released insulin crystal.

## Introduction

Examples of over 50 biologically active peptides/low molecular-weight substances released by exocytosis include body’s only glucose-lowering hormone insulin, adrenaline regulating the ‘flight-or-fight’ response as well as dopamine, serotonin and large neurotransmitter molecules(Golan et al., 2011). Our understanding of secretory (patho)physiology is increasingly based on real-time functional studies, a powerful tool that allowed pin-pointing functional defects observed in such diseases as diabetes mellitus(Collins et al., 2016). For instance, advances in high resolution fluorescence live imaging, particularly super-resolution stimulated emission depletion (STED), allowed real-time visualisation of a fusion pore dynamics thereby providing a better understanding of secretion mechanisms(Shin et al., 2018; Zhao et al., 2016). However, as any nanoscale fluorescent reporter is likely to affect the secretory kinetics, every molecular detail of insulin release that we are aware of comes from surrogate molecules(Gandasi et al., 2018; Low et al., 2014; Tarasov et al., 2018) or indirect measurements(Hastoy et al., 2018; Li et al., 2011). The need for revealing the secretory machinery as it happens in real life suggests for less invasive live-imaging techniques capable of direct, label-free visualisation of DCV release, which could also be combined with fluorescence imaging.

Biophysical alternatives to fluorescence microscopy of single-vesicle exocytosis are nominally free from the label problem; at the same time, they are less informative than imaging and, again, do not measure insulin release directly. Monitoring of single-vesicle exocytosis using patch-clamp electrophysiology(Neher & Marty, 1982) or electrochemical techniques(Huang et al., 1995; Zong et al., 2010) provides excellent signal-to-noise ratio but largely lacks spatial information. This problem may be resolved by using scanning probe microscopy (SPM) techniques. Atomic force microscopy (AFM) has been applied to imaging of regulated exocytosis in AT2 cells but the prohibitively high forces applied by AFM probe keep the resolution of the technique way off the individual DCV range(Hecht et al., 2012). Scanning Ion Conductance Microscopy (SICM) has been successfully utilised for the detection of the exocytotic sites by the scanning probe in chromaffin cells, fixed before and after the stimulation(Shin & Gillis, 2006).

Our approach, a combination of optical microscopy and SPM techniques, is based on a simultaneous acquisition of the label fluorescence and rasterising the ‘surface landscape’ of the cell membrane. The fluorescent label has an auxiliary role assisting the selection of the region of interest (ROI) for the scanning probe, a glass nanopipette that follows the surface of a sample within the ROI very closely albeit without touching. In this way, correlative scanning ion conductance microscopy - fluorescence confocal microscopy (SICM-FCM) imaging effectively operates directly and non-invasively, which makes it a perfect fit for live-cell studies. A recent improvement of the acquisition rate has allowed us to monitor secretion of individual insulin containing vesicles in real time.

## Methods

### SICM-FCM

#### Principles of operation

SICM generates high-resolution (down to ~5nm) topographical images of samples immersed in liquid by raster scanning a glass nanopipette (probe) that follows the surface of a sample very closely without touching (Adenle & Fitzgerald, 2005; Hansma et al., 1989; Korchev et al., 1997; Shevchuk et al., 2006). Importantly, SICM sets no requirement for any kind of chemical processing of the studied sample and effectively operates in native conditions. This fully non-invasive aspect of imaging makes SICM ideal for living cell studies. The SICM instrument (Fig. 1A) consists of a nanopipette mounted on a Z piezo actuator connected to a scan control unit that performs vertical measurements and raster-scanning XY piezo actuator that carries the cell sample in a Petri dish. Control electronics measures the ion current that flows between the probing and the reference electrodes located inside the SICM pipette and in the bath, respectively. When the pipette approaches the sample surface to approximately one radius of its [pipette tip] opening, the ion current drops (Fig. 1B). This drop is used by the control software to stop the approach and record the vertical position of Z piezo actuator as the sample height at this imaging point. The pipette is then withdrawn, and sample is moved to the next location (Fig. 1A, red arrows). Such regime of scanning is termed Hopping Probe SICM (HPICM)(Novak et al., 2009).

**Fig. 1.**
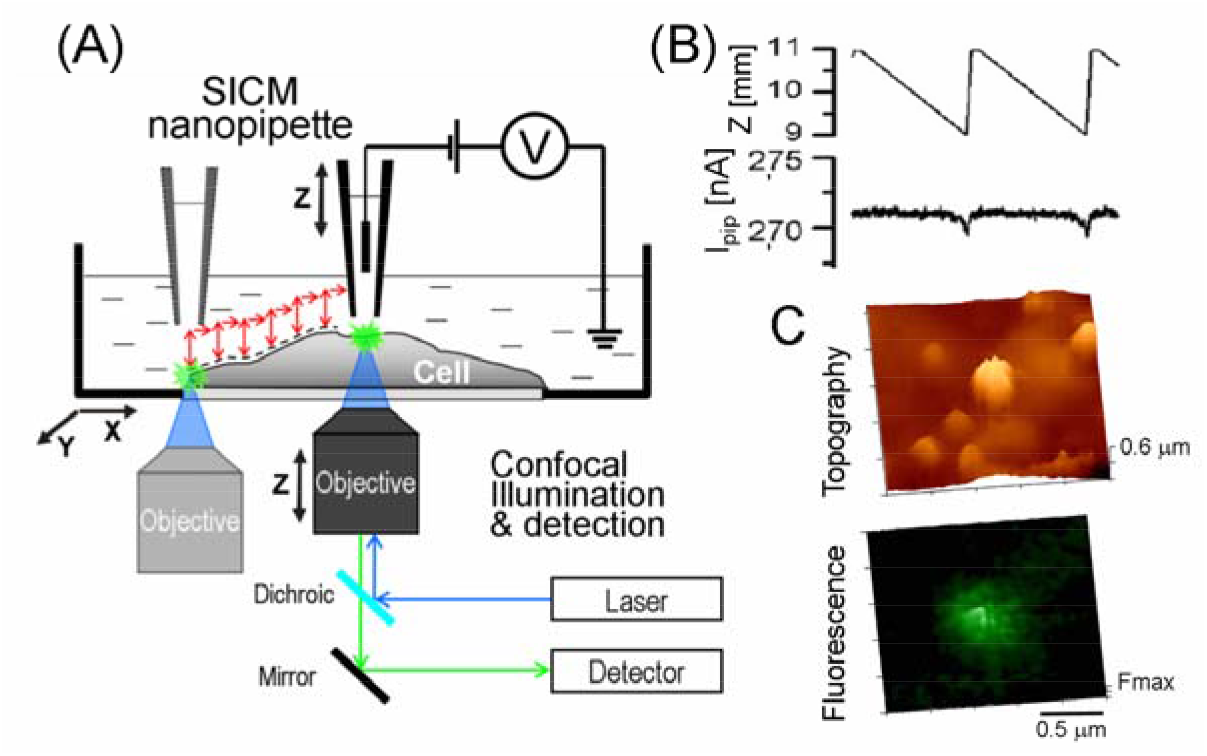
SICM-FCM principles of operation. **(A)** Schematic diagram of simultaneous, correlative scanning ion conductance and fluorescence confocal microscope. **(B)** Vertical position (Z) and corresponding ion current (I_pip_) traces during two adjacent points measurement. **(C)** SICM topography and confocal image pair shows insulin granule on the INS-1E cell plasma membrane and corresponding NPY-Venus fluorescence.

Simultaneous fluorescence confocal imaging is performed by a laser beam focused at the pipette tip (Gorelik et al., 2002). This provides co-localized fluorescence excitation, which is then recorded by a photomultiplier. Confocal auto-focusing is achieved by vertical movement of the objective focusing piezo actuator that is synchronised with the SICM height measurement Z piezo actuator. This eliminates the need for 3-D reconstruction through numerous confocal slices and allows the fluorescence data to be acquired precisely from the apical membrane in single scan (dashed line, Fig. 1A). Figure 1C shows an example of a 3-D topographical and matching fluorescence images of an insulin granule releasing at the apical surface of an INS-1E cell and corresponding NPY-Venus fluorescence targeted to the granules by means of fusion with neuropeptide Y. Like other scanning probe microscopy techniques, SICM image generation relies on the scanning of its probe over the sample surface. Since piezo actuators used to move the probe have finite time response, the imaging rate depends on the image size and the number of pixels comprising the image. It has been demonstrated that it is possible to design and construct SICM that could be operated in high speed (HS) regime sufficient to follow highly dynamic processes in living cells (Ida et al., 2017; Simeonov & Schäffer, 2019a, 2019b).

All experiments were performed using a custom built SICM setup consisting of XY-stage (45 x 45 μm, Direct Drive, Capacitive Sensors, Parallel Metrology, IC-XY-4545-001) and Z-stage (25 μm, Direct Metrology, Capacitive Sensor, IC-Z-25-001) powered by Piezo Controller System (IC-PDC-001) and operated using ICAPPIC Controller (IC-UN-001, ICAPPIC Ltd., UK). To allow high speed imaging the setup was equipped with a fast Z piezo actuator (IC-HSZ-001, ICAPPIC Ltd. UK). Fine alignment of the scanning nanopipette tip with the laser beam in the XY plane was done using two N-470 PiezoMike Linear Actuators (Physik Instrumente, Germany) and was based on the optical image acquired using a 100× oil immersion objective. Therefore, partial offset of fluorescence images in relation to topography could be observed. Images were superimposed by adjusting fluorescence image contrast to saturation and tracing the edges of fluorescence spots using ImageJ software (https://imagej.nih.gov/ij/). Then the background of fluorescence images was set transparent and the images overlaid on corresponding topographical image.

Nanopipettes were pulled from BF-100-50-7.5 borosilicate glass capillaries (Sutter Instrument Co., USA) using a P-2000 laser puller (Sutter Instrument Co., USA). Ion currents were measured using an Axopatch 200B amplifier (Molecular Devices, UK) with a gain of 1 mV/pA and a low-pass filter setting of 5 kHz. The internal holding voltage source of the Axopatch 200B was used to supply a direct current voltage of +200 mV to the pipette. The ion current and outputs of the capacitive sensors from all three piezo elements were monitored using Axon Digidata 1322A and Clampex 9.2 (Molecular Devices, UK).

Fluorescence images were recorded using a D-104 Microscope Photometer (Photon Technology International, Inc. USA) through a 100×/1.3NA oil immersion objective. The excitation was provided by a 488-nm wavelength diode-pumped solid-state laser (Laser 200; Protera) focused to diffraction limited spot. Confocal volume height was 1.5 μm estimated by imaging of 100nm TetraSpeck microspheres (T7279, Life Technologies Corporation, Oregon, USA) in XZ plane.

SICM-FCM control, data acquisition and analysis were performed using HPICM Scanner and SICM Image Viewer software (ICAPPIC Ltd, UK) custom modified by authors.

### Cell culture and plasmids

INS1-E cells were cultured in RPMI1640 medium (11 mM glucose) supplemented with 10% of fetal bovine serum and 50 μM of β-mercaptoethanol. Cells were cultured under sterile conditions, 37°C, 100% humidity. Prior to the experiments, INS1 cells or primary β-cells were plated on glass-bottomed dishes (MatTek Life Sciences, USA) and NPY-Venus (kindly provided by Atsushi Miyawaki, RIKEN BSI, Wako, Japan) was delivered adenovirally, with 24-36-h expression time.

### Animals and islet isolation

C57Bl/6 mice (Charles River, Harlow, UK) were used throughout the study. Mice were kept in a conventional vivarium with a 12-hour-dark/12-hour-light cycle and ad libitum access to food and water and were killed by cervical dislocation. All mouse experiments were conducted in accordance with the United Kingdom Animals (Scientific Procedures) Act (1986) and the University of Oxford ethical guidelines.

Pancreatic islets were isolated from mice by injecting liberase (Type V, Sigma) solution into the bile duct, with subsequent digestion of the connective and exocrine pancreatic tissue. Islets were picked using a P20 pipette, under a dissection microscope and cultured overnight in RPMI medium containing 11mM glucose, supplemented with 10% FBS, 100IU/mL penicillin and 100μg/mL streptomycin (all reagents from Life Technologies/ThermoFisher, Paisley, UK). Islets were dispersed into single cells by pipetting in low-Ca^2+^ extracellular medium.

## Results

### Morphological changes associated with insulin secretion in INS1-E cells

We first studied insulin DCV release in the rat insulinoma cell line, INS-1E. SICM topographical images of control uninfected INS1-E cells revealed the cell surface decorated with numerous microvilli and dorsal ruffles that could be seen as membrane protrusions (Fig. 2A). We then imaged INS1-E cells infected with a type 5 human adenovirus delivering a fluorescent label of insulin DCVs, NPY-Venus(Nagai et al., 2002; Tarasov et al., 2018). NPY is a 36 amino-acid neuropeptide that is known to be packaged into insulin containing granules and is widely used as an insulin secretion marker. SICM-FCM images of non-stimulated cells revealed similar microvillar and dorsal ruffle structures (Fig. 2B, black and white arrows). Superimposing fluorescence and topographical images (see Methods) revealed smooth membrane bulges spatially correlating with NPY-Venus fluorescence spots, which could be interpreted as membrane-docked insulin-containing vesicles (Fig. 2B, red arrows). The depth of the confocal volume (~1.5 μm) allows detecting the NPY-Venus fluorescence signal from not yet docked insulin vesicles located beneath the membrane. This explains why the majority of fluorescent spots did not co-localise with protruding membrane structures, in non-stimulated cells.

**Fig. 2.**
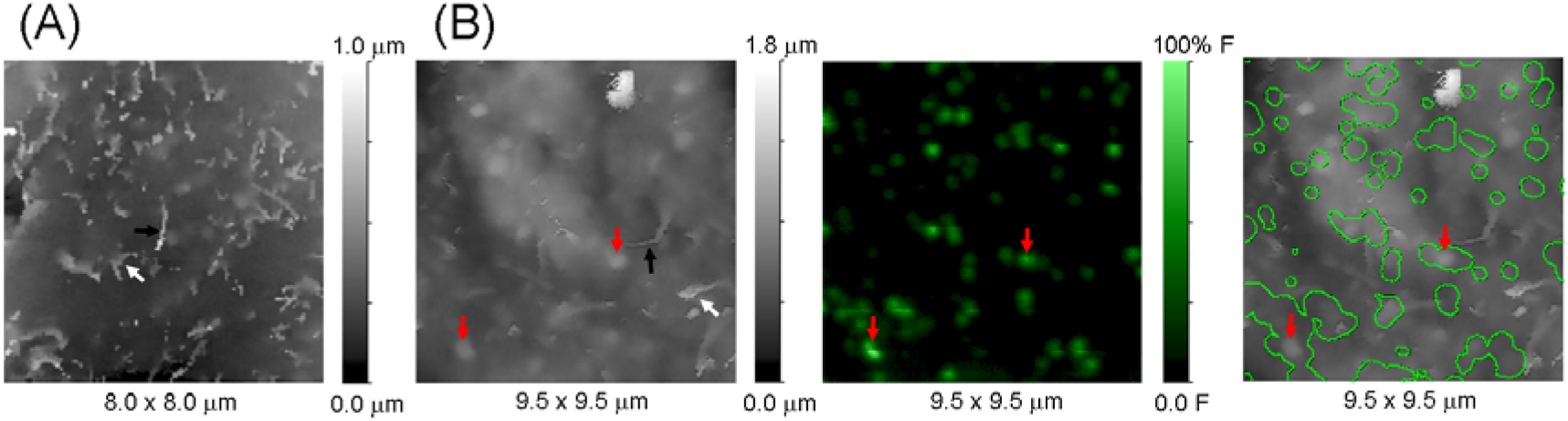
INS1-E cell surface morphology. **(A)** Typical SICM topographical image of control (uninfected) INS1-E cell showing microvilli (black arrow) and dorsal ruffles (white arrow). **(B)** Correlative SICM (left panel), fluorescence confocal (middle panel) and overlay (right panel) images of non-stimulated INS1-E cell expressing NPY-Venus fixed on ice showing microvilli (black arrow), dorsal ruffles (white arrow) and membrane-docked insulin DCVs (red arrows).

To estimate the density and frequency of the fusion events, we imaged INS1-E cells expressing NPY-Venus fixed 10 and 30 seconds after stimulation with 17 mM glucose and 0.2 mM tolbutamide (Fig. 3A and B, respectively). Numerous hemispherical protrusions with diameters ranging from 200 to 400 nm were detectable, 10 s after stimulation (Fig. 3A),. While some protrusions matched NPY-Venus fluorescence (red arrow), representing vesicles fixed at the early stage of release, several protrusions without any associated fluorescence signal (cyan arrow) were also seen.

**Fig. 3.**
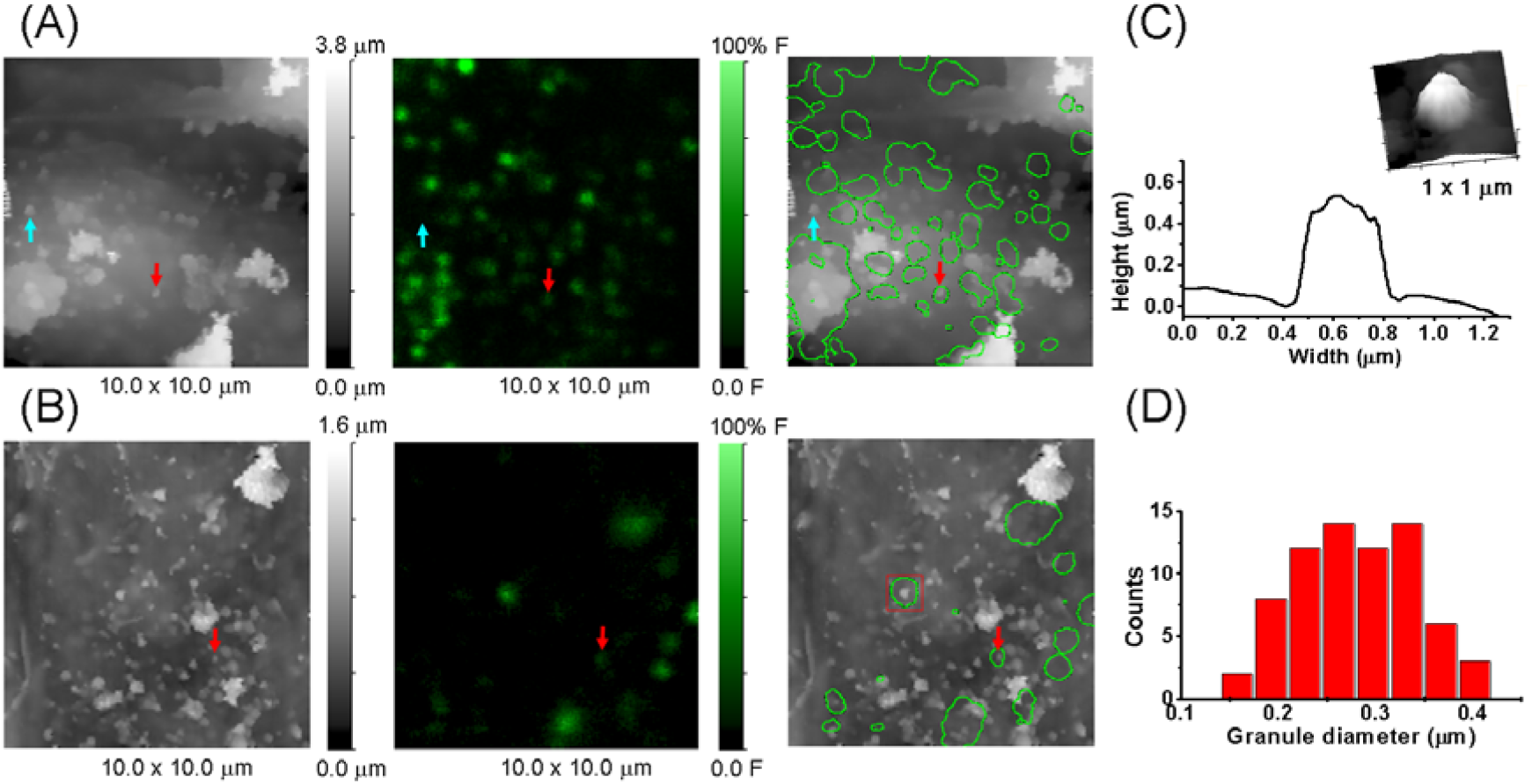
Correlative SICM-FCM images of insulin granules on surfaces of stimulated INS1-E cells expressing NPY-Venus. **(A)** Images of INS1-E cell fixed 10 seconds after stimulation showing released insulin granules with (red arrow) and without (cyan arrow) associated NPY-Venus fluorescence. **(B)** Images of INS1-E cells fixed 30 seconds after stimulation showing released insulin granules. Few granules have associated NPY-Venus fluorescence (red arrow). **(C)** Cross section profile and zoomed 3D topographical image (inset) of typical released insulin granule boxed in (B). **(D)** Distribution of insulin granule diameters measured by SICM.

SICM-FCM images of cells fixed 30 seconds after the stimulation show larger numbers of insulin granules on the cell membrane and fewer NPY-Venus fluorescence maxima (Fig. 3B), as compared to cells fixed 10 seconds after stimulation (Fig. 3A). Yet, we still were able to detect several fluorescent spots correlating with protrusions (red arrow in Fig. 3B). A typical protrusion had linear size of 200-400 nm and ~500 nm height (Fig. 3C, representing the zoomed red box in Fig. 3B). The distribution of protrusion size measured by SICM had the mean diameter of 0.28±0.06μm (n=71) (Fig. 3D), in reasonable agreement with values for insulin crystals of 150-200 nm, estimated by TEM (Raleigh et al., 2017). We therefore conclude that, correlative SICM-FCM imaging can identify and topographically resolve individual insulin granules on the cell membrane of the INS1E cells.

### Kinetics of insulin dense core vesicle release in INS-1E cells

To resolve the kinetics of DCV fusion and subsequent insulin granule release, we performed correlative time-lapse SICM-FCM imaging of stimulated INS1-E cells (Fig. 4). We observed a NPY-Venus fluorescence maxima marking a spot of subsequent topographically detected protrusion corresponding to released insulin granules (Fig. 4A, first frame, labelled ‘00:00’). This corresponds to the docking of the insulin-loaded vesicle to the cytoplasmic side of the cell membrane(Whim, 2011). A rapid appearance of a protrusion at the spot (cyan arrow) is likely to reflect the vesicle fusion and exposure of the insulin granule on the cell surface (Fig. 4A, 3^rd^ frame, ‘00:36’, indicated by red arrow). From this moment, both the size of the protrusion and the intensity of NPY-Venus fluorescence begin to drop, suggesting the dissolution of the insulin crystal (Fig. 4B, frames 4-7, ‘00:54’ onwards). We also observed events in which protrusions that correlated spatially with NPY-Venus fluorescence spot appeared only in one frame. These could represent insulin granules that detach from the cell membrane. The average granule release/dissolution time measured from the moment of its appearance on the cell membrane to complete disappearance was 40±21 sec. (n=7). Reported by light microscopy, the diameter of 20-μm synthetic insulin crystals reaches the limit of detection (~200 nm, corresponding to a 100-fold decrease in the linear size) between 210 and 440 seconds, depending on the presence/absence of albumin in the solvent (Lougheed et al., 1981). Likewise, we earlier reported a several minute-long delay between the peak post-secretory endocytosis and bulk insulin levels in the extracellular medium (Tarasov et al., 2018). A tenfold decrease of the protrusion diameter, from ~300 nm (Fig. 3A) down to 30 nm, a small size particle that could be resolved with confidence by SICM on a surface of living cell, fits very well within this range.

**Fig. 4.**
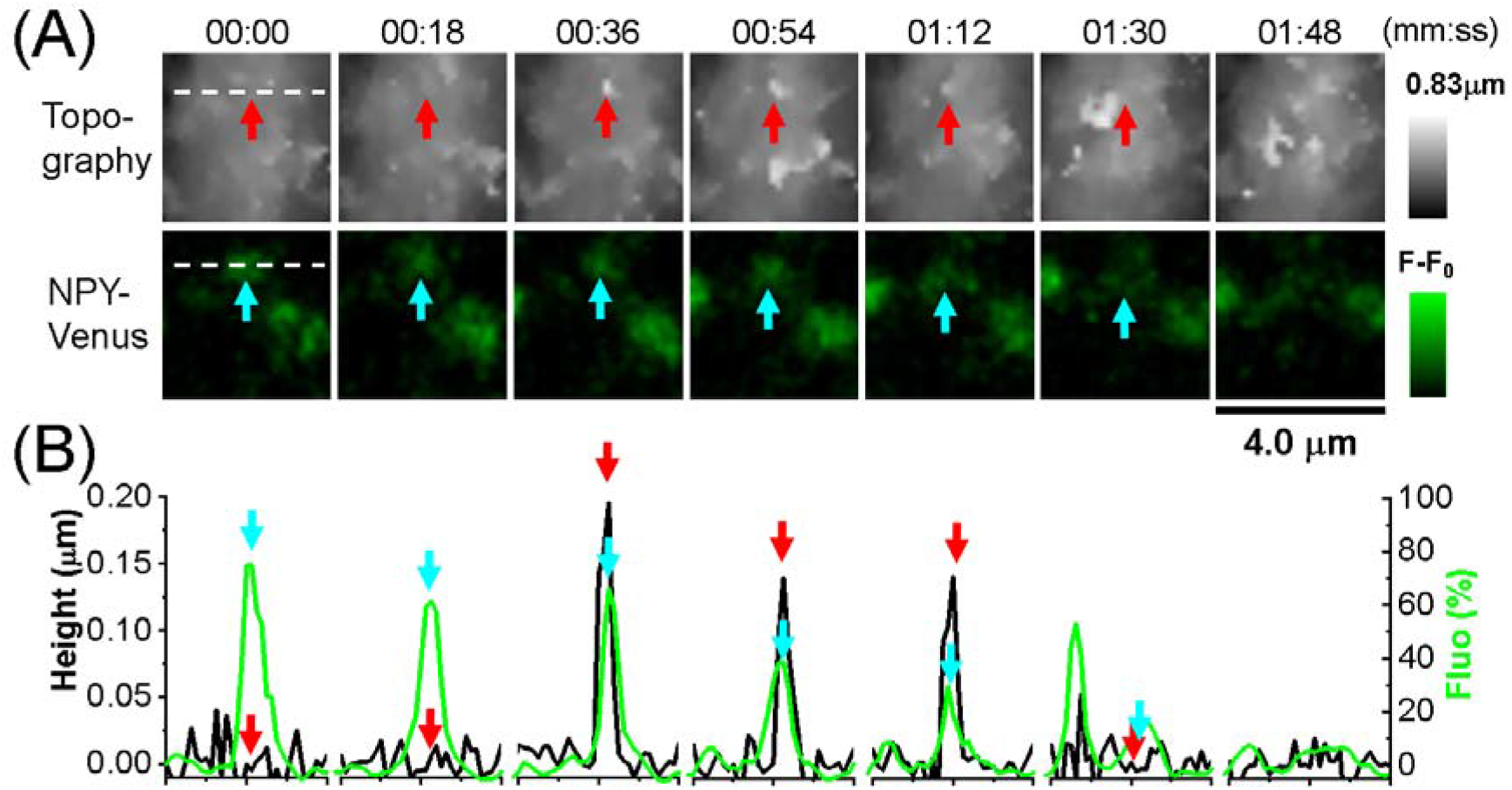
Insulin DCV secretion in INS1-E cells expressing NPY-Venus. **(A)** Topographical (top row) and NPY-Venus fluorescence (bottom row) time lapse images showing appearance of insulin granule on the cell surface (red arrow at 00:36 seconds) and corresponding NPY-Venus fluorescence signal (cyan arrow) recorded ~2 min after stimulation. **(B)** Cross section profiles of SICM topography (black trace) and fluorescence intensity (green trace) for each frame at location marked with white dashed line in the first frame (A).

A possible cause for the delay between the dissipation of the NPY-Venus fluorescence and the protrusion reported at the putative site of exocytosis is the difference in kinetics of the dissolution of the crystal and diffusion of a small protein off the cell surface. Differently from the inverted TIRF imaging, both freshly released NPY-Venus and insulin crystal are facing a much larger volume of the medium, hence both processes are likely to proceed faster, on the top, unrestricted surfaces of cells.

On several occasions, we were able to detect an indentation in the cell membrane that preceded the arrival of the NPY-labelled insulin granule to the cell membrane (Fig. 5A, red arrow, third frame at 36 seconds). Based on the spatial and temporal correlation of topographically detected insulin granule and NPY-Venus fluorescence spot we conclude the indentation represents the full opening of the fusion pore, perhaps reflecting the final stages of full fusion of the insulin vesicle with the plasma membrane. The pore opening was 370 nm measured as full width at half minimum that is in agreement with recently published STED observations (Shin et al., 2018).

**Fig. 5.**
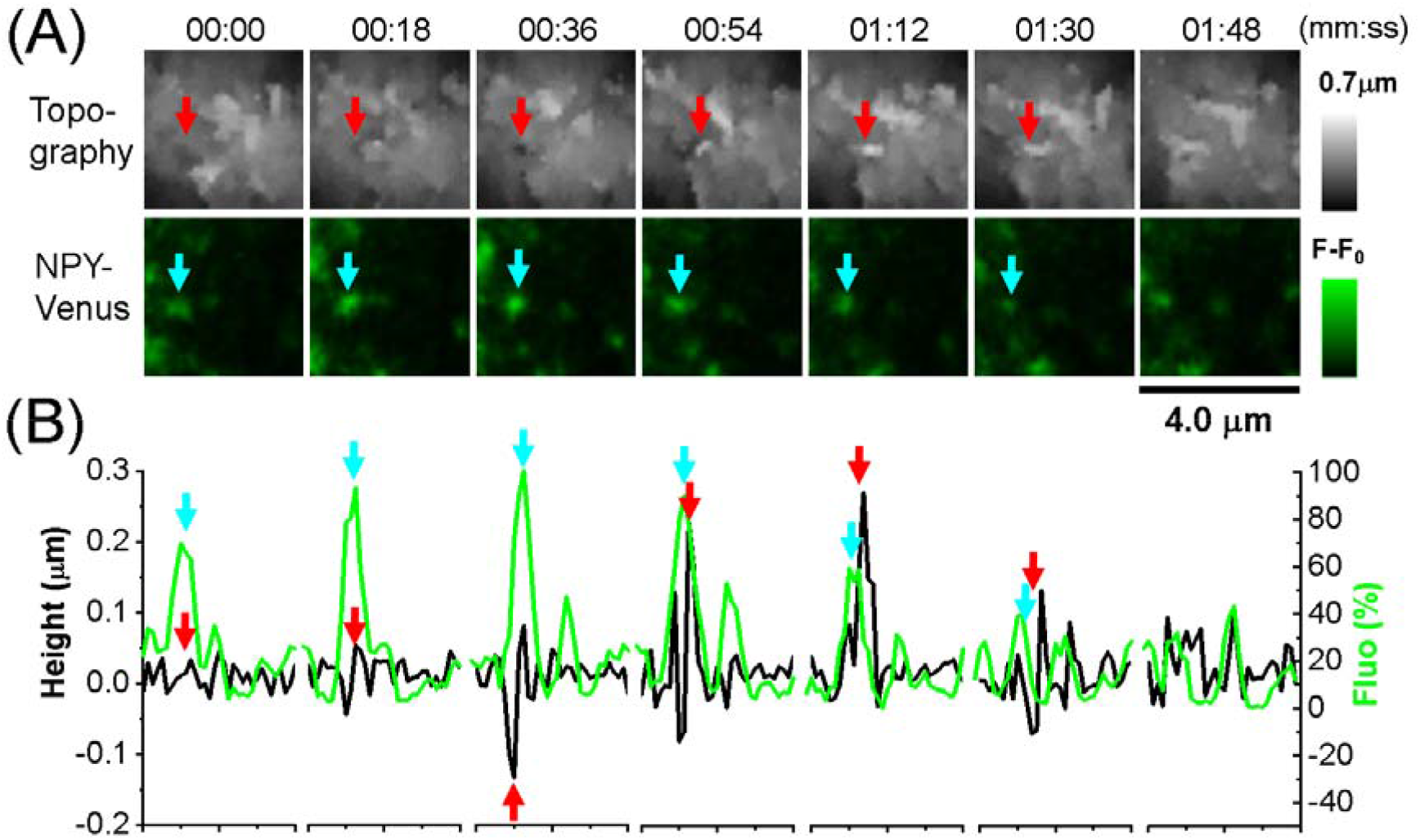
Opening of the fusion pore during complete fusion. **(A)** Topographical (top row) and NPY-Venus fluorescence (bottom row) time lapse images showing the opening of the fusion pore (red arrow at 00:36 seconds) followed by the appearance of insulin granule on the cell surface (red arrow at 00:54 seconds), and corresponding NPY-Venus fluorescence signal (cyan arrow) recorded ~2 min after stimulation. **(B)** Cross section profiles of SICM topography (black trace) and fluorescence intensity (green trace) for each frame at location marked with white dashed line in the first frame (A).

### Insulin dense core vesicle release in primary β-cells

We then applied the imaging technology to primary mouse pancreatic islets and imaged the secretion of unlabelled insulin granules in primary mouse β-cells, with the stimulation being provided locally via a micropipette (Fig. 6A). The stimulation resulted in drastic cell surface changes that could be observed in SICM topographical images (Fig. 6 B). Similar to what we observed in INS1 cells, the protrusions resembling insulin granules by size and shape, within several minutes after the stimulation of β-cells (Fig. 6C). The average release and dissolution time of putative insulin granules in β-cells was 52±25 sec (n=3), close to that observed for INS1-E cells.

**Fig. 6.**
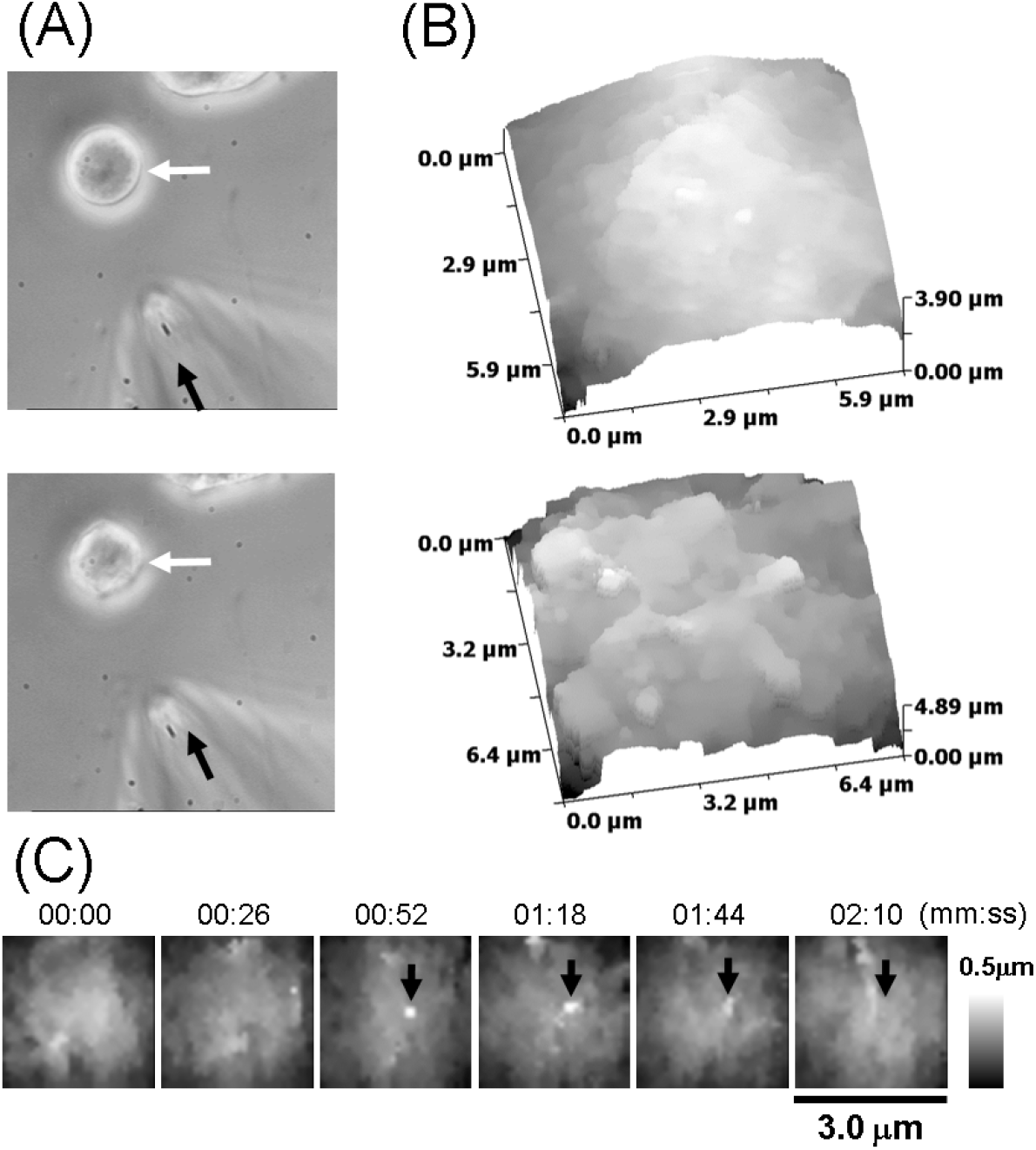
Insulin DCV secretion in primary b-cells. **(A)** Bright filed images of a β-cell (white arrow) before (top panel) and after (bottom panel) localised stimulation through micropipette (black arrow). **(B)** SICM topographical images of a β-cell before (top) and after (bottom) stimulation. **(C)** Series of SICM topographical time lapse images showing the release and dissolution of insulin granule (red arrow) recorded ~4 min after stimulation.

## Discussion

Although hormone DCV release was studied in detail using fluorescence live imaging of associated reporter molecules, little is known about the secretion and dissolution kinetics of individual granules of insulin itself. Insulin is densely packed within the β-cell DCV, with the reported concentrations of 118 and 74 mM for human and rat β-cells, respectively(Rorsman & Renström, 2003). The peptide has been shown to exist in a form of heterohexamer (MW=36 kDa, R_h_=5.5-5.6 nm(Hvidt, 1991; Oliva et al., 2000)) at concentrations exceeding 2 mM(Hansen, 1991). Insulin is therefore likely to be stored within the vesicles predominantly as a crystallised hexamer(Brange & Langkjœr, 1993), possibly accompanied by a small soluble fraction. The latter point has been long supported by the electron microscopy (EM) of chemically fixed cells reporting insulin secretory granules (ISG) as 200-mn electron-dense cores, surrounded by a 100nm lighter halo, both encapsulated within a lipid membrane(Dean, 1973; Sorra et al., 2006). The authenticity of the halo, conventionally believed to contain soluble insulin, was, however, recently been questioned: the halo was not found in high-pressure frozen preparations and therefore was referred as a chemical fixation artefact(Fava et al., 2012). A similar halo was reported in chromaffin cells, with the ability to detect the halo depending on the fixation method(Stevens et al., 2011).

Based on monitoring of fluorescent insulin surrogates (typically, GFP variants fused to DCV membrane proteins or cargo molecules)(Tsuboi & Rutter, 2003) or Zn^2+^ (Li et al., 2011; Michael et al., 2006; Qian et al., 2003) (an ion that stabilises insulin hexamer, can be visualised with a number of specific dyes) using total internal reflection fluorescence (TIRF) microscopy, two distinct modalities have been suggested for the insulin vesicle exocytosis. The release of insulin via a transient fusion pore, via the so-called ‘kiss-an-run’ process avoiding the full fusion of the vesicle and liberation of its nonsoluble contents, was suggested based on monitoring NPY-Venus kinetics(Tsuboi & Rutter, 2003). The full fusion was believed to represent a relatively slower (1-2s) and irreversible event resulting in the dissolution of the insulin crystal in extracellular milieu(Michael et al., 2006).

The weakness of these fluorescence measurements is that the size of the reporter is different to that of insulin and so is likely to be its kinetics. The fluorescent tag (YFP) fused to vesicular cargo (NPY) is a protein with molecular weight over 26.9 kDa, and R_h_ of 2.2 nm (Hink et al., 2000) i.e. less than a half of the radius of the vesicular hexameric insulin (5.6 nm(Hvidt, 1991)), perhaps more capable of reflecting the dynamics of insulin monomers or dimers (2.7 and 3.8 nm, respectively(Oliva et al., 2000)).

The observed release kinetics of the chimaeric construct might therefore represent that of the tag rather than the cargo protein itself (Michael et al., 2006), which becomes critical, when considering the ability to pass through the fusion pore of a size of 1.4-1.6 nm(Kasai et al., 2010; MacDonald et al., 2006). Thus, the true ability of hexameric insulin to leave through the fusion pore remains questionable. Complicating matters, in technical sense, TIRF microscopy does not report fusion of vesicles with extracellular medium; instead it monitors vesicles fusing with a small confined 30-50nm space between the glass and the cell membrane. This is highly likely to affect the rate of a 250-nm (Hink et al., 2000) crystalline insulin core release and dissolution as the volume-limited diffusion certainly enhances the observed spatial and temporal correlation between insulin and fluorescent markers.

In our hands, the released insulin granule had a measurable lifetime of 40±21 s at the cell membrane, correlating well with earlier observations of insulin crystal dissolution (Lougheed et al., 1981) as well as high-content imaging of endocytosis in islets (Tarasov et al., 2018). This finding can also explain a ~2-min delay between the depolarisation-induced influx of Ca^2+^ and insulin secreted by pancreatic islets and detected in a soluble form in the bath medium (Ravier et al., 1999).

## Conclusion

Here we show that high speed correlative SICM-FCM can be used to visualise both unlabelled insulin granule and fluorescently labelled insulin DCV release from the top, non-adherent surface of primary pancreatic β-cells as well as β-cell lines, such as INS1-E cells. These results establish correlative SICM-FCS as a method of choice for studying the molecular mechanisms of diabetes induced by impaired or deficient hormone secretion from pancreatic islet cells as well as profiling novel drugs targeting pancreatic hormone release.

## Abbreviations

MW: molecular weight
R_h_: hydrodynamic radius

## Acknowledgements

This work was supported by BBSRC funding to A.S. (BB/M022080/1). P.N. acknowledges support from the Russian Science Foundation (grant no. 19-79-30062). Y.K. acknowledges support from the World Premier International Research Center Initiative (WPI), MEXT, Japan.

